# A map of human protein-protein interaction embeddings for functional discovery

**DOI:** 10.64898/2026.08.07.743440

**Authors:** Mert Cihan, Ute Distler, Miguel A. Andrade-Navarro

## Abstract

A protein’s function depends not just on its own structure and localization, but also on the interactions with its partners. Many proteins are therefore better described by a set of partner-dependent roles than by a single annotation. Yet most approaches to the functional interpretation of protein-protein interactions (PPIs) remain protein or set-centric. They rely on pre-existing annotations, and perform worst where knowledge is sparse. Here, we present MAPPIE (Map of Protein-Protein Interaction Embeddings), a method that treats each PPI, rather than each protein, as a unit of representation. From 199,137 human interactions spanning 15,503 proteins, we build a two-dimensional map of the human PPI landscape for functional discovery. Protein language model embeddings for two protein interaction partners are combined and compressed into a latent space, with model selection guided by domain-domain interactions used as a structural proxy for interaction similarity. The resulting geometry separates domain defined interaction classes, organizes disorder associated interactions spatially, and splits interactions involving the same protein by partner. A query PPI’s latent neighbourhood recovers its own annotated functions across molecular, complex, pathway, and biological processes. MAPPIE contributes most where existing functional evidence is weakest, outperforming interactome and sequence identity baselines for sparsely connected interactions. MAPPIE neighbours of query PPIs are enriched for partners in independent protein networks, recovering curated complex-level function even when subunits are spread across the map. Applied to a human dark interactome, MAPPIE assigns specific, experimentally supported functions to dark hub proteins.

## Introduction

Protein function is highly context dependent. Although amino-acid sequence constrains the structural and biochemical potential of a protein, its realized role in the cell is shaped by abundance, subcellular localization, and the molecular partners it engages (1, 2). Protein–protein interactions (PPIs) underlie many cellular processes by organizing proteins into functional complexes and regulatory networks. These interactions can be stable or transient, constitutive or context-specific, and may involve structured domains or short recognition elements in disordered regions (3, 4). A protein’s biological role is therefore often better understood as a set of partner-dependent functions than as a single intrinsic annotation. This is especially true for hub proteins, where distinct edges in a protein interaction network correspond to distinct molecular consequences (1, 5).

Large-scale experimental and curated resources have substantially expanded the known human protein interactome, establishing protein connectivity as a central layer linking genotype to phenotype (6, 7). Yet functional interpretation of PPIs still largely relies on protein- or set-centric strategies. The interactors of a bait protein, or the members of a network cluster, are tested for whether Gene Ontology, pathway, or domain terms are overrepresented, or graph proximity is used to prioritize genes and functions at scale (6, 8). These approaches are necessary, yet they share a fundamental limitation. Because they depend on pre-existing annotation, they perform worst where knowledge is most sparse, and they characterize neighborhoods rather than the mechanistic consequence of any single interaction. Moreover, they collapse heterogeneous relationships into a single measure, which may reflect direct binding, co-complex membership, pathway adjacency, co-expression, or literature mention (9).

For an individual PPI, biological interpretation ideally operates at finer resolution, capturing not only whether two proteins interact, but how, where the interface is encoded, and what context makes the interaction meaningful. Domain–domain interaction catalogs, short linear motif resources, and disorder-focused databases each provide a partial route toward this goal (10, 11), and structure prediction has recently made complex-level modeling far more accessible (12). Still, these methods do not solve *de novo* functional interpretation at interactome scale. A structural model may suggest interface geometry but not pathway role, regulatory consequence, or how an interaction is positioned relative to thousands of related PPIs. What is missing is a representation in which interactions themselves, rather than proteins alone, are the primary computational objects, and in which partner-specific context becomes visible as geometry.

Protein language models (PLMs) provide an attractive basis for such a representation because they shift part of the functional signal from external curation back to sequence. Trained by self-supervised learning on large bodies of protein sequences, PLMs generate dense embeddings that capture biochemical, evolutionary, and structural regularities without task-specific labels, and have supported prediction of structure itself directly from sequence (13, 14). Critically, every protein, including poorly characterized or sparsely connected ones, can be assigned a comparable representation independent of interactome coverage. PLMs are therefore a natural foundation for representing interactions, yet existing sequence-based PPI methods use them primarily to predict whether two proteins interact, rather than to represent interactions as objects whose arrangement encodes partner-specific function (15).

Here we describe MAPPIE (Map of Protein-Protein Interaction Embeddings), a method for interaction-level functional prediction in a PLM-derived embedding space. Rather than limiting analysis to individual proteins, MAPPIE treats each PPI as a distinct interaction-level representation. PPIs are encoded from PLM-derived protein embeddings, combined into interaction-level vectors, compressed into a latent space, and projected onto a fixed two-dimensional manifold. The same protein can therefore populate different regions of the map when paired with different interactors, making partner-specific context explicit. The reference set comprises 199,137 human PPIs spanning 15,503 proteins. A webserver supports protein search and highlighting, inspection of individual interactions, and projection of user-supplied PPIs from UniProt identifiers or FASTA sequences. Moreover, MAPPIE allows for functional term discovery through enrichment analysis over Gene Ontology (GO) terms (16, 17), KEGG (18) and Reactome pathways (19), PFAM (20) and InterPro (21) domains, domain-domain interactions (DDIs) from 3did (22), and biochemical reactions from Rhea (23), for a PPI or set of PPIs through neighbourhood retrieval. The resulting MAPPIE geometry reveals multiple layers of organization, including separation of domain-defined interaction classes, stronger local purity for disorder-associated programs, and partner-specific splitting of interactions involving the same protein. MAPPIE is freely available at https://cbdm-01.zdv.uni-mainz.de/∼mcihan/mappie/.

## Results

### Domain-domain interfaces structure the MAPPIE landscape

Creating a map of PPIs through compression of combined protein language model embeddings is an unsupervised task that requires no prior knowledge beyond the interacting partners and their sequences. However, to evaluate which of the generated maps is most representative of a functional landscape, without making that information explicit during map construction, we leverage interface-centric information by annotating DDI interfaces onto the mapped PPIs (Fig. S1). We reasoned that if two proteins interact through a pair of domains documented to physically bind, their interaction is structurally well-supported, and a representative map should place such interactions close together.

We addressed this by scoring each DDI with at least two mapped PPIs by the compactness of its interactions in latent space, the mean distance of its PPIs to their centroid, normalized by the global mean across DDIs. Across all tested combinations of embedding source (ESM-2 and ProtBERT), merge operation, and latent dimensionality, this identified ESM-2 (13) with 128 dimensions and elementwise multiplication as one of the top performers, achieving a score of 0.61 (Fig. S2). This combination remained top-ranked across different embedding sources and confidence thresholds. Our primary reference set comprises the 199,137 HIPPIE (7) interactions retained at a confidence score ≥ 0.64. Restricting instead to a higher-confidence subset (the top 10% of HIPPIE entries, score ≥ 0.82) further improved the score from 0.61 to 0.70, but we retained the primary set to preserve coverage of all 199,137 interactions.

To characterize how DDIs are organized in MAPPIE, we computed a local purity score relative to the background rate (Fig. 1A). We found that 286 DDIs with more than 30 annotated PPIs show a fold-enrichment of median 173-fold over the random baseline (Fig. 1B), confirming that structurally-related interactions are locally clustered.

**Fig. 1.**
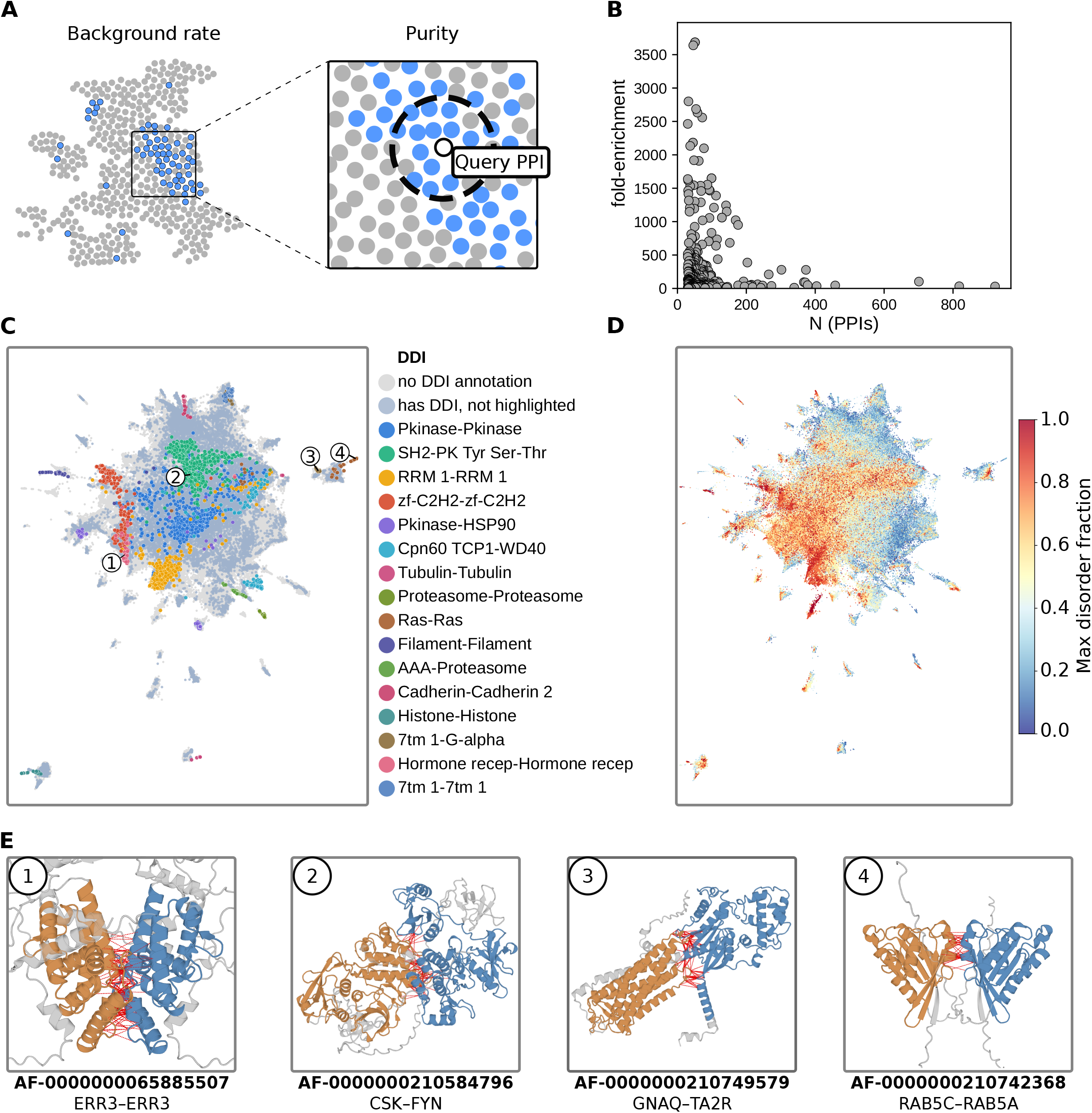
Structure guided selection of latent dimension. **A**) Schematic representation showing local enrichment of DDI-annotated PPIs around a query interaction relative to their background. PPIs sharing the query’s DDI are shown in blue and all remaining (background) PPIs in grey. **B**) Fold-enrichment of each DDI with >30 PPIs annotated within its local neighbourhood (*k* = 15) plotted against the number of PPIs carrying that DDI. The baseline is each DDIs background frequency in the full PPI set. **C**) PPIs colored for selected DDIs in MAPPIE. **D**) MAPPIE coloured by the maximum disorder fraction of the two interacting proteins. **E**) AlphaFold-Multimer structural models of the four example PPIs highlighted in **C**). The two Pfam domains of the interacting proteins are coloured orange and blue, and inter-chain residue contacts within 10 Å are shown as red lines.

We make this visible by coloring selected homo- and heterodimeric DDIs in MAPPIE. Interaction classes such as Pkinase–Pkinase, RRM 1–RRM 1, and zf-C2H2–zf-C2H2 occupy larger but separated territories, while others such as 7tm 1–G-alpha and Pkinase–HSP90 form small, well-separated islands at the periphery (Fig. 1C). These selections can be performed and explored using the web server.

Colouring the map by the maximum disorder fraction of the two partners shows that proteins with high disorder fraction also organize spatially, forming separated regions rather than spreading uniformly across MAPPIE (Fig. 1D).

Moreover, for four example PPIs drawn from distinct DDI clusters (Fig. 1C, circles 1–4) with high-confidence AlphaFold-Multimer predictions available (24), we confirmed that the annotated domains genuinely form the interface. We find inter-chain contacts within 10 Å (142 contacts for ERR3–ERR3, 84 for CSK–FYN, 90 for GNAQ–TA2R, 42 for RAB5C–RAB5A) fall predominantly between the two annotated domains (orange and blue, Figure 1E). This supports the interpretation that co-localization on the map may reflect shared interface architecture.

### Function is recoverable across scales

There is no single definition for the function of a PPI. The same PPI may be described by the interaction itself, molecular activity, complex membership, pathway, or the broader biological process it serves. Rather than committing to one definition of a biological function, we asked whether MAPPIE’s latent organisation recovers annotated terms across all of these scales at once, spanning molecular activity, complex membership, pathway, and processes. If interactions of a kind lie near one another on the map, a query’s own functions should be recoverable from its neighbourhood.

Term annotation coverage was near-complete for the InterPro, GO CC, Pfam, GO BP, and GO MF categories, each assigned to over 98% of PPIs in MAPPIE. Pathway and reaction sources covered fewer interactions, down to RHEA (38.3%) and structural information for DDIs are the sparsest (13.7%; Figure 2A). The number of GO biological processes dominates at 4,944,383 PPI–term associations (Fig. 2B).

**Fig. 2.**
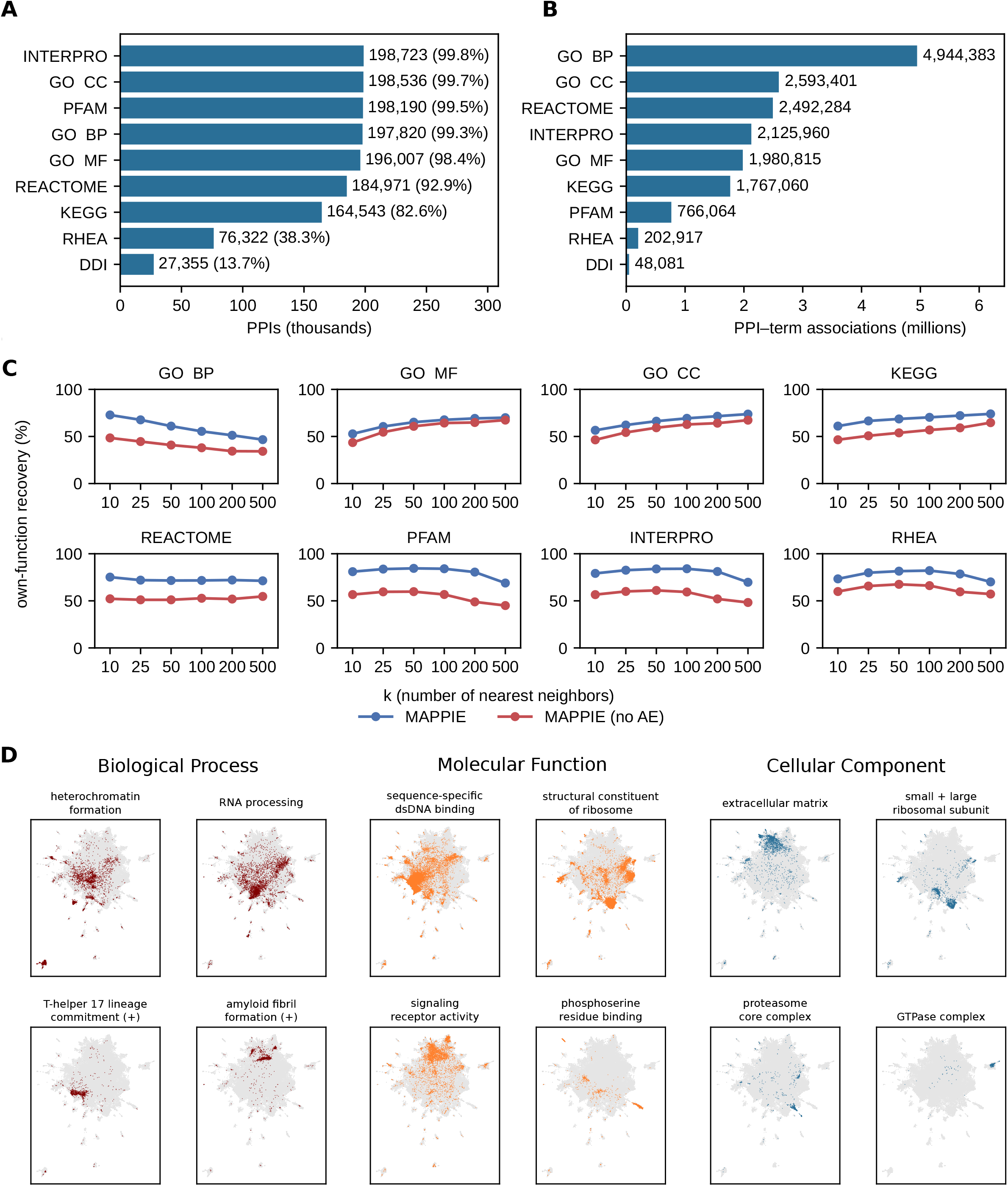
Functional annotation and function recovery. **A**) Number of MAPPIE PPIs annotated by each functional category. **B**) Total number of PPI–term associations per category. **C**) Own-function recovery as a function of neighbourhood size for each annotation category, comparing the full MAPPIE model against a control using the interaction embedding without autoencoder compression. **D**) PPIs annotated with example GO Biological Process, Molecular Function, and Cellular Component terms colored in MAPPIE.

To evaluate performance across all categories we then tested whether a query PPIs latent neighbourhood recovers its own annotated functions (Fig. 2C). The GO biological process recovered best in the tightest neighbourhood (72.8% at k = 10) and declined as the neighborhood grew, indicating BP terms are concentrated locally and diluted by more distant interactions. GO MF, GO CC and KEGG instead improved with k, reaching 70.0%, 73.8%, and 74.0% respectively at k = 500, consistent with these functions spanning broader regions. The domain categories and RHEA terms peaked at intermediate sizes (k ≈ 50–100) before falling.

In every category, the full model outperformed the control lacking autoencoder (AE) compression (Fig. 2C, blue vs. red). Compressing the multiplied PLM embeddings into a compact latent space therefore does not merely preserve but sharpens the functional neighbourhood, recovering a query’s own functions more reliably. This advantage persists when excluding neighbours that share either partner with the query, ruling out partner reuse as the source (Fig. S3). The margin is smaller across the GO branches, whose recovery relies more on partner-sharing neighbours than the other categories. We therefore conclude that MAPPIE places PPIs together based not only shared function, but also on sequence similarity.

We highlight that individual terms capture spatially separated regions in MAPPIE rather than scattering across the map uniformly (Fig. 2D). Broad terms such as sequence-specific double-stranded DNA binding (n = 11,669) and extracellular matrix (n = 2,208) span large but coherent areas, while specific terms such as GTPase complex (n = 260) and T-helper 17 cell lineage commitment (n = 398) concentrate in small, sharply-localised clusters (Fig. 2D). This holds across all three GO branches, indicating the map organizes interactions by function across scales.

### Function recovery beyond homology and interactome baselines

Functional inference for proteins is conventionally derived from interactome-centric annotation transfer, where a protein inherits the functions of its network neighbours through the “guilt by association” method (9, 25), and from sequence similarity to already-characterized proteins (16, 26). We asked whether the MAPPIE neighbourhood recovers PPI functions beyond these homology and network-based inference methods.

For each query PPI, we compared four sources of candidate proteins matched to the same size: the first-degree interactome neighbours from HIPPIE, MAPPIEs latent neighbours, MAPPIEs combined embedding without AE compression, and a sequence-identity baseline. We stratified queries by network degree to test how each method behaves as prior knowledge of an interaction grows (Fig. 3A). To make the comparison strict, we excluded any candidate sharing 30% or more sequence identity with the query and restricted GO annotations to experimentally supported terms (see Methods).

**Fig. 3.**
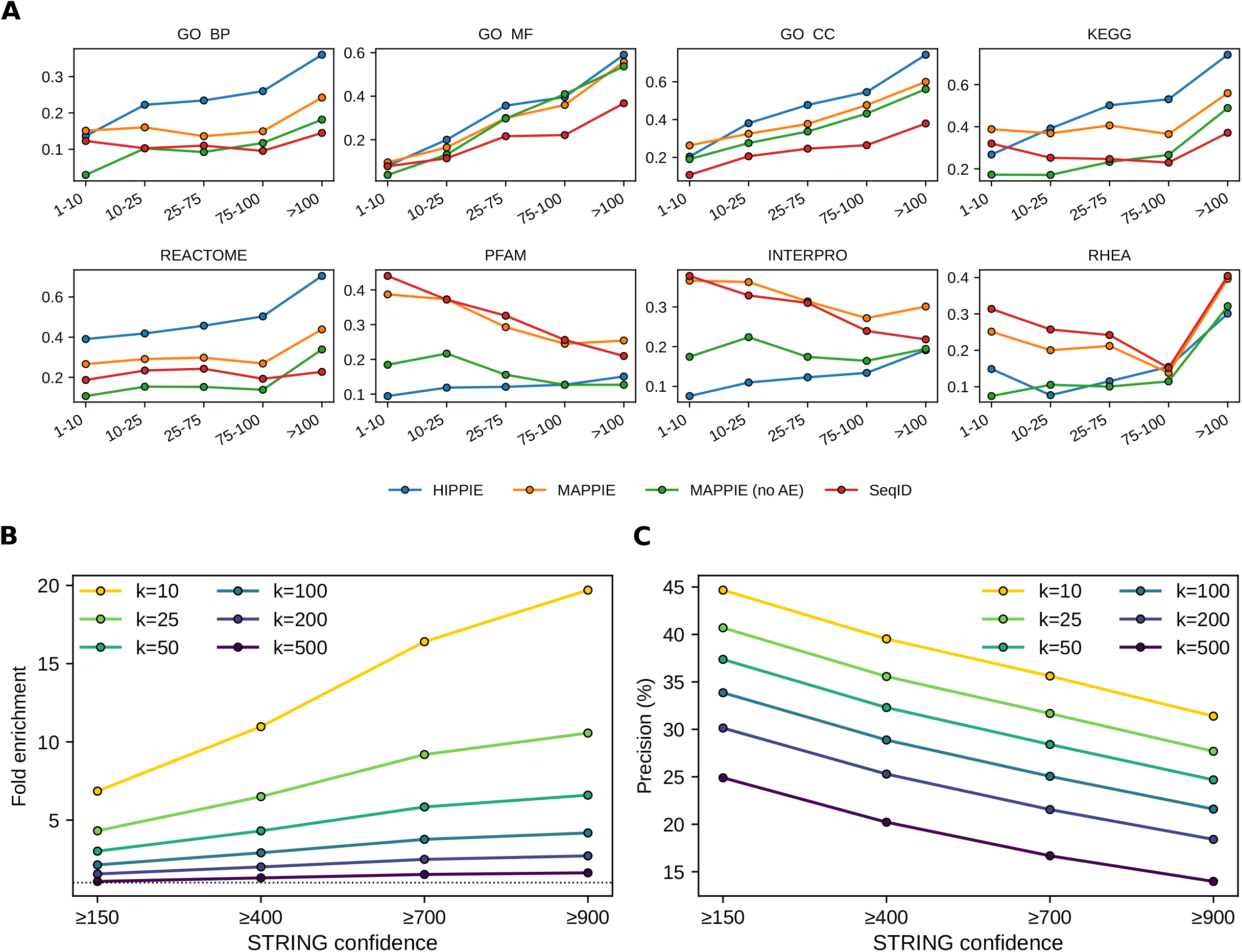
Benchmarking against baselines and an external network. **A**) Own-function recovery for each annotation category, stratified by network degree of a query PPI, comparing MAPPIE against network first-degree neighbourhoods, the combined interaction embedding without autoencoder, and a sequence-identity baseline. **B**) Fold enrichment of STRING network recovery for STRING confidence thresholds, shown across neighbourhood sizes. **C**) Precision of STRING network recovery across the same confidence thresholds and neighbourhood sizes.

For sparsely-connected interactions, MAPPIE recovered functions better than the interactome baseline across the GO categories and KEGG pathways (GO BP: 15.1% vs 13.7%; GO MF: 9.6% vs 8.1%; GO CC: 26.3% vs 20.4%; KEGG: 38.8% vs 26.8%; at 1–10 partners) (Fig. 3A). As the number of known partners grew beyond ten, this advantage reversed. Here, HIPPIE’s network-based selection has an increasingly informative neighbourhood to draw on and became the strongest method, forming an effective ceiling for the process- and pathway-level categories at the highest degrees. The observation therefore is that MAPPIE contributes the most where existing interaction evidence is weakest, and the biophysical interactome baseline dominates only where the query’s interactome is already densely characterized.

MAPPIE also performed consistently better than its no-AE compression control across these stratifications, confirming the effect stems from the compressed latent organisation rather than the combined protein language model embedding alone. For the domain categories the sequence-identity baseline was strongest for PPIs with few partners (Pfam: 44.0%; InterPro: 37.8%; at 1–10 partners), despite the exclusion of close homologues, as expected since domains are encoded in sequence. Recovery dropped with degree for both, since domains cluster tightly in MAPPIE and wider neighbourhoods weaken the signal (Fig. 3A).

To validate against an independent network, we asked whether proteins close in MAPPIEs latent space are also connected in STRING. For each MAPPIE PPI in STRING, we counted how many of its nearest latent neighbours corresponded to STRING edges, relative to a degree-corrected random expectation (Fig. 3B; see Methods). MAPPIE neighbours were more likely to be STRING partners than chance, growing with STRINGs PPI confidence score. At k = 10, enrichment rose from 6.9× (≥ 150) to 19.7× (≥ 900) with precision reversing this trend (Fig. 3C), being higher for tighter neighbourhoods (37.8% at k = 10 vs 18.9% at k = 500). Proximity in MAPPIE therefore enriches in true protein interactions, confirmed by an independent network.

### Complex-level function recovered from multiple clusters

Because MAPPIE places each PPI independently, the PPIs of a given protein complex can sit in different regions of the map depending on its partners. We asked whether these scattered interactions still recover the complex’s curated function. For each human complex from CORUM, we reconstructed the PPIs among its subunits, pooled the 100 latent neighbours of each into a single set, and tested that combined neighbourhood for enrichment of the complex’s annotated GO terms (see Methods).

Across 302 human complexes with two or more mappable PPIs, MAPPIE recovered the curated function in most cases, with a mean GO recall of 72.0% (Fig. 4A). The terms for Spliceosome (100%) and Acetylcholine receptor signaling pathway (100%) were recovered completely, while Regulation of oxygen-sensing pathways (45%) and Visual perception (50%) sat lower.

**Fig. 4.**
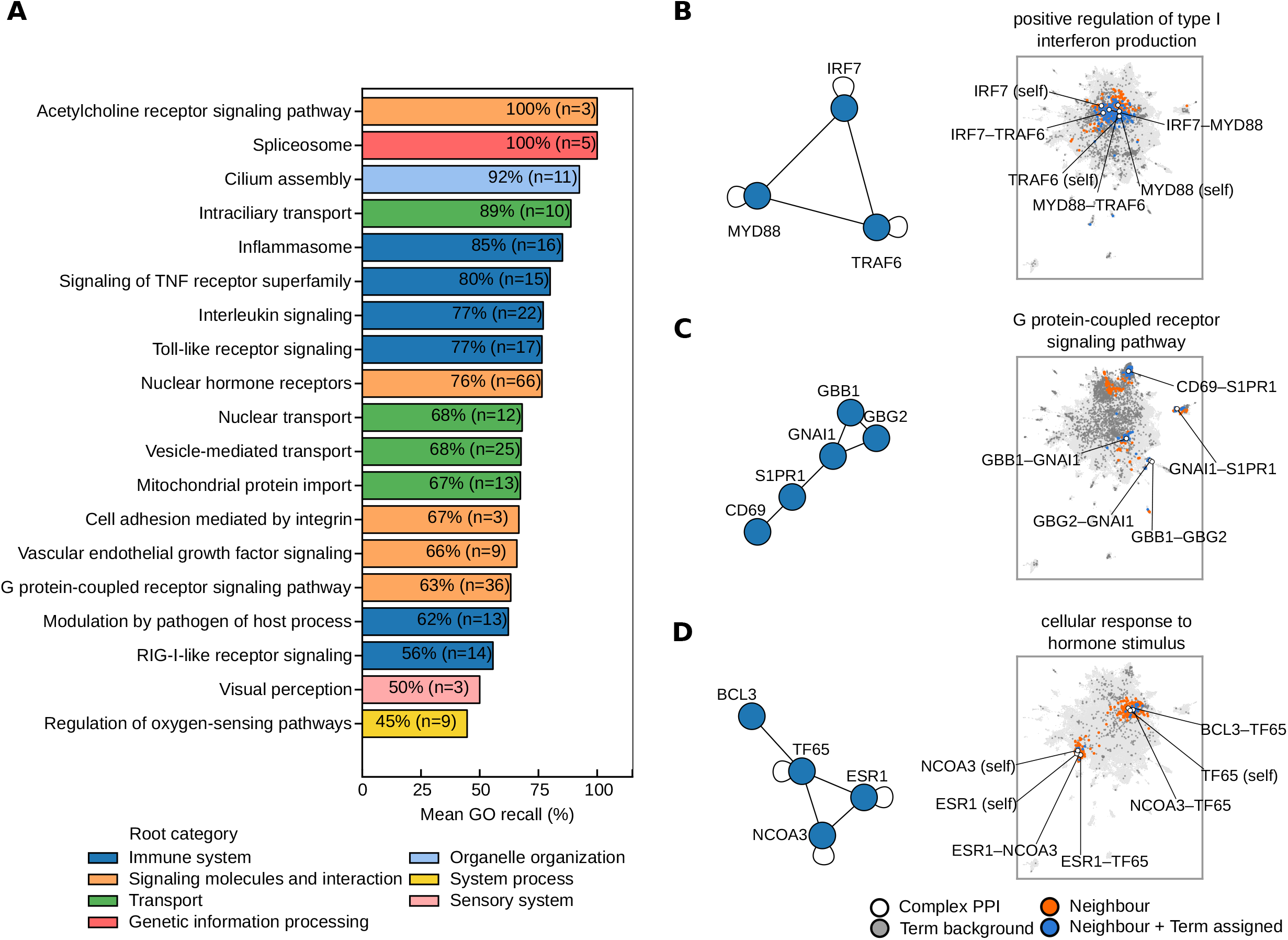
Recovery of curated protein complexes. **A**) Mean GO recall for protein complexes, grouped by their curated functional category and coloured by root category. **B–D**) Three example complexes, each with its protein interaction subgraph on the left and its annotated PPIs on the MAPPIE, shown with their latent neighbours. Neighbours that also carry the complex’s GO term against the terms overall background across the map are highlighted. The recovered GO term is given above each panel: positive regulation of type I interferon production **B**), G protein-coupled receptor signaling pathway **C**), and cellular response to hormone stimulus **D**).

To illustrate the term enrichment across a complex distributed over several PPIs, we highlight three examples (Fig. 4B–D). In each, the subunit interaction subgraph (left) maps onto a set of PPIs that individually sit among neighbours carrying the shared term (right). For the type I interferon complex (IRF7–MYD88–TRAF6; Figure 4B), the constituent PPIs cluster together in one region, and each sits within a neighbourhood enriched for positive regulation of type I interferon production, so the term is recovered locally for every PPI (191/473 pooled neighbours carry this term; adj p < 0.05). The G-protein-coupled receptor example (Fig. 4C) is more dispersed: the core G-protein PPIs (GBB1–GNAI1, GBG2–GNAI1, GBB1–GBG2) group in two regions while the receptor-facing interactions (CD69–S1PR1, GNAI1–S1PR1) sit in separate parts of the map, yet all regions are enriched for G protein-coupled receptor signaling pathway (277/414 pooled neighbours carry this term; adj p < 0.05). For the hormone-response complex (ESR1–NCOA3–TF65–BCL3; Figure 4D), the interactions occupy neighbouring but distinct positions, each embedded among PPIs annotated to cellular response to hormone stimulus (80/510 pooled neighbours carry this term; adj p < 0.05).

Together these show that MAPPIE recovers complex-level function without requiring a complex to form a single cluster, since each of its PPIs is independently embedded among functionally similar interactions.

### Functions of the human dark interactome

Many proteins in the human proteome remain functionally uncharacterized, yet a substantial number of them have experimentally supported interaction partners. We defined a human dark interactome of 1,925 PPIs by taking high-confidence BioPlex interactions (pInt ≥ 0.75) (5) and keeping only those in which both partners are classified as functionally dark (knownness score ≤ 1.0 in the Unknome database (27)), then computed the functional enrichment with MAPPIE. Within this network, proteins had on average 3.2 interactions (Fig. 5A). For each of these PPIs we retrieved its latent neighbourhood and predicted GO term enrichment (see Methods), recovering on average 96.8, 26.9 and 24.6 significantly enriched terms for biological process, molecular function and cellular component, respectively.

**Fig. 5.**
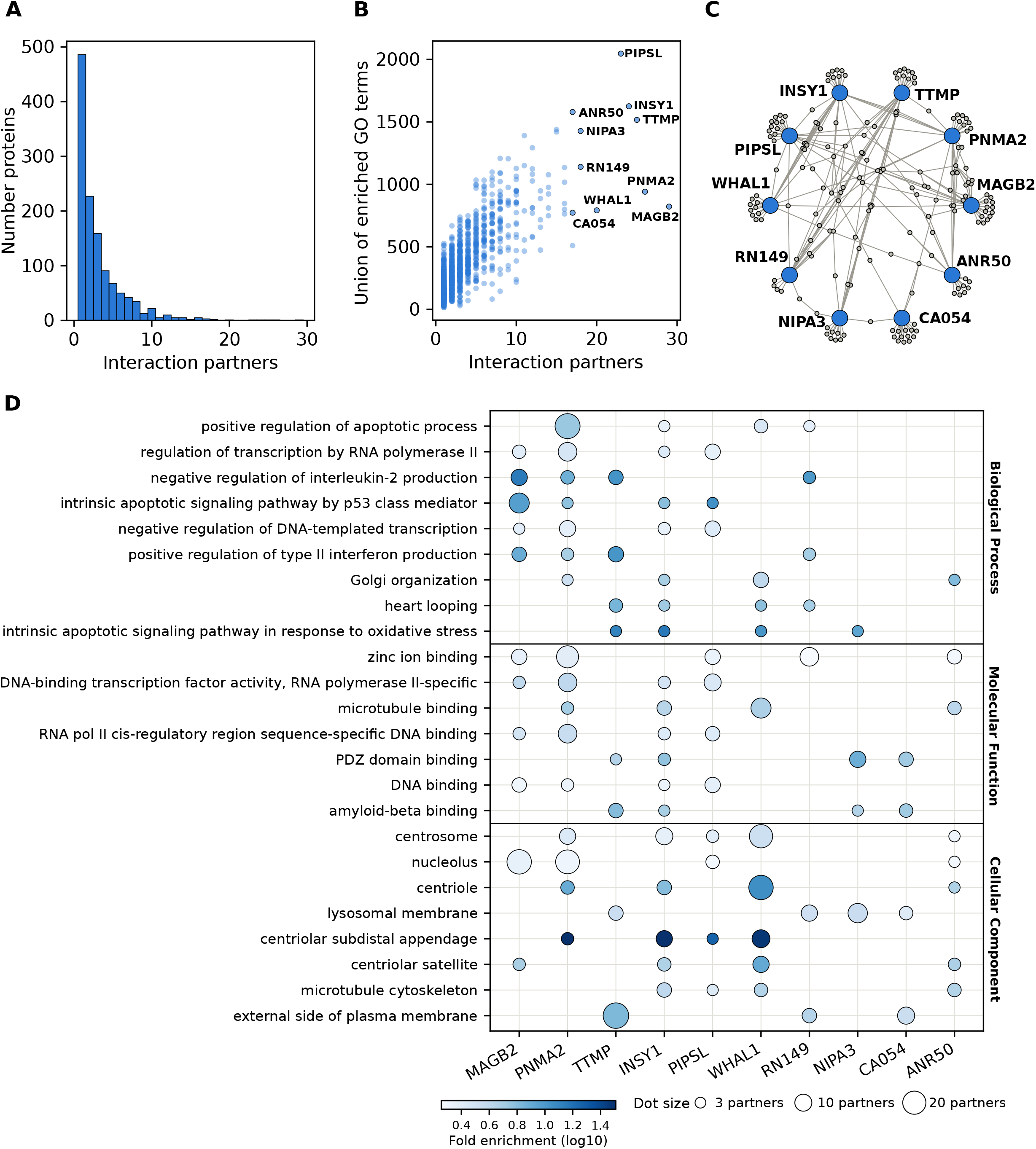
Hub proteins of the dark interactome. **A**) Distribution of the number of PPIs per dark protein. **B**) Degree versus the union of GO terms significantly enriched (hypergeometric test, BH-adjusted p ≤ 0.05) across all of a protein’s dark PPIs. The ten most-connected hub proteins are labelled. **C**) Interaction subgraph of the same ten hub proteins from B) and their direct dark PPIs. **D**) Recurrent GO terms enriched across the dark PPI neighbourhoods of the ten hub proteins (≥ 3 PPIs supporting the same term per hub).

For the ten most-connected hub proteins in this network we find on average 1,266 enriched terms in their PPIs (Fig. 5B); their interaction subgraph is shown in Figure 5C. Together these hubs and their PPIs cover 217 interactions representing 11.3% of the dark interactome. We reasoned that when the same function is enriched across the neighbourhoods of many PPIs, this function can be attributed to the interaction properties of the hub protein itself, rather than to any single PPI. We highlight terms enriched across ≥3 PPIs of a hub as candidate annotations for that dark hub protein (Fig. 5D).

The functions recovered for these hubs extended beyond broad, high-level ontology terms. Within biological process, PNMA2 was enriched for positive regulation of apoptotic process across 21 of its 26 PPIs, and MAGB2 for the more specific intrinsic apoptotic signalling pathway by p53 class mediator across 13 of its 29 PPIs. Interestingly, both of these apoptotic predictions are supported by recent, independent experimental work. The knockdown of PNMA2 directly promotes apoptosis in endometriotic cells (28) while depletion of MAGB2 increased PARP cleavage, consistent with enhanced apoptosis, in medulloblastoma cells (29) (Fig. 5D).

RNF149 was enriched for interferon-related regulation across 6 of its 18 PPIs, consistent with recent work showing that it ubiquitinates IRF3 to drive its proteasomal degradation and suppress interferon production (30) (Fig. 5D).

Together, these examples show that MAPPIE can assign specific, experimentally supported functions to proteins that carry no usable annotation of their own and would be invisible to conventional annotation transfer. We provide the full list of GO annotations for the dark interactome in Dataset S1.

## Discussion

In this work we introduced MAPPIE, a map of human protein–protein interaction embeddings in which functionally related interactions occupy neighbouring regions of a shared latent space. By treating each PPI, rather than each protein, as the unit of representation, MAPPIE makes partner-specific context explicit as geometry. This reflects a broader principle that a protein’s role is often better described as a set of partner-dependent functions than as a single intrinsic annotation, particularly for hub proteins whose distinct edges carry distinct molecular consequences (1, 5).

Our benchmarks show that the annotated functions of a PPI are recoverable from its latent neighbourhood across molecular, complex, pathway, and process scales, and that this holds even for interactions whose proteins were excluded from close sequence homologues. Not every term is recovered for every interaction, which we attribute to the context-specificity of function itself. A single protein contributes different roles to different complexes, so no neighbourhood is expected to capture the union of a protein’s annotations, but rather those relevant to the specific pairing. Such context-dependent multifunctionality is well documented in moonlighting proteins, where a single polypeptide performs distinct functions that can depend on its interaction partners and cellular context (31, 32).

Combining protein embeddings into an interaction-level object for functional discovery, rather than for binary interaction prediction, distinguishes MAPPIE from existing PLM-based approaches that study PPIs. These approaches resolve interactions at finer structural detail, yet they characterize how and if two proteins meet rather than what the interaction does (15, 33, 34). MAPPIE instead asks what an interaction potentially means at interactome scale. It does so without end-to-end fine-tuning of PLMs, but is based on a frozen protein language model (13) and a lightweight compression model, which keeps the method simple and easy to extend to new proteins or protein fragments.

The principal strength of this design is that functional inference becomes sequence-based. Conventional annotation transfer relies on interactome membership through *guilt-by-association* (9, 25) or on sequence similarity to characterized proteins (16, 26), both of which fail where prior knowledge is sparse. The query in MAPPIE does not need to have prior annotation or prior interactome membership. In principle this allows any candidate pair including engineered or mutated sequences, or protein fragments, to be projected and interpreted. Consistent with this, MAPPIE contributed most where existing evidence was weakest, complementing rather than replacing interactome-based inference. We illustrate this on the human dark interactome, where interactions between proteins that lack usable annotation of their own still recover specific functions, candidate apoptotic roles for the hub proteins PNMA2 and MAGB2 (28, 29), and interferon regulation for RNF149 (30) each independently supported by recent experimental work.

Several limitations follow from the same design. First, the reference interactome is incomplete, with high-throughput mapping estimated to cover well under a quarter of all pairwise human interactions (35), so absence from a region reflects sampling as much as biology. Second, MAPPIE places any submitted pair on the map regardless of whether the two proteins genuinely interact. It is a tool for functional interpretation, not interaction validation, and remains complementary to dedicated interaction predictors and structural modelling (12, 15). Third, functional recovery is bounded by the completeness of the underlying annotation, so sparsely annotated regions are evaluated conservatively and may harbour genuine functions we cannot yet score. Moreover, because most available functional annotations are protein-level, the benchmark uses the union of the two partners annotations and therefore does not by itself establish that every recovered term is specific to the interaction rather than to one of its interacting proteins.

Future work could replace the frozen, mean-pooled representation with interaction-aware embeddings that explicitly model the interface, for instance the pairwise residue representations learned by interface-centric models (33), which we expect to sharpen the separation of interaction classes further. Extending the map beyond human proteomes and incorporating conditional context such as tissue or subcellular localization (2) are natural further directions.

By making interactions themselves the objects of a functional landscape, MAPPIE offers a method to interpreting the specific functions of individual interactions across a large fraction of the interactome that previously received only a single, pooled functional annotation.

## Materials and Methods

### MAPPIE architecture overview

MAPPIE embeds PPIs in a shared latent space in which functionally related interactions lie close together, enabling map-based visualisation, nearest-neighbour retrieval, and functional annotation transfer. MAPPIE is based on PLM embeddings that encode each protein, a merging step that reduces the two partner embeddings to a single vector, and an autoencoder (AE) that compresses this vector into a compact latent representation used for PPIs projection. To select representative parameters for the final map, domain–domain interactions (DDIs) are mapped to PPIs and used as a structural proxy to score how tightly related interactions cluster (Fig. S1).

### Protein-protein interactions

PPIs for MAPPIE were taken from HIPPIE v2.3 (7), which assigns confidence scores by integrating evidence from multiple experimental sources weighted by the number and type of supporting experiments. We applied a score threshold of ≥ 0.64 (covering 27% of scoring interactions), resulting in 199,137 PPIs for 15,503 unique proteins. For comparison, we additionally defined another high-confidence subset comprising the top 10% of HIPPIE entries (corresponding to a score threshold of ≥ 0.82), resulting in 80,893 PPIs for 12,605 unique proteins (Fig. S4).

### Pair representation

For each PPI, we encoded both proteins with a pretrained PLM, using ESM-2 (650M) (13) as the primary source and generating ProtBERT (36) embeddings for comparison. We mean-pooled the final-layer per-residue representations, giving each protein a single fixed-length vector of 1280 dimensions for ESM-2 and 1024 dimensions for ProtBERT. We then reduced each protein pair to a single interaction vector using one of four merge operations, tested independently: the elementwise average, the element wise multiplication, the absolute difference, and the ordered concatenation in both possible directions, which doubles the dimensionality.

### Latent compression

We standardized each interaction vector to zero mean and unit variance and compressed it with an autoencoder (AE) implemented in PyTorch (37). The encoder is a 4-layer MLP with fixed intermediate widths of 512 and 256 and a swappable bottleneck (the final encoder layer) of 512, 256, or 128. The decoder mirrors this, with ReLU activations throughout and linear final layers, reconstructing the input under mean squared error. Input dimensionality is set by embedding source and merge operator: 1280 (ESM-2) or 1024 (ProtBERT) for average, multiply, and difference, and double that for concatenation. Each network was trained for 20 epochs (Adam, learning rate 1e-3, batch size 1024) on a 90/10 train/validation split, retaining the lowest-validation-loss epoch. We repeated this across the full grid of embedding source (ESM-2, ProtBERT), merge method (average, multiply, difference, concatenation), bottleneck dimensionality (128, 256, 512), and HIPPIE (7) confidence filter (≥ 0.64; ≥ 0.82) (Fig. S2).

### Model selection

To select a representative map of PPIs for functional discovery, we map domain–domain interactions (DDIs) from 3did (22) using Pfam (20) annotation to all PPIs in MAPPIE and leverage them as a structural proxy for interaction similarity. PPIs sharing a documented DDI should lie close together. In total, 6,374 DDIs mapped to PPIs (48,081 DDI–PPI instances). Restricting to the 4,364 DDIs present under both HIPPIE confidence filters, we scored each DDI *d* mapping to at least two PPIs by the compactness of its PPI set *P(d)*, the mean distance of its PPIs to their centroid *c(d)*:

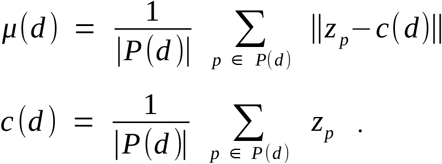

where *z*_*p*_ is the latent embedding of PPI *p*. We normalized each DDI’s compactness by the global mean across DDIs:

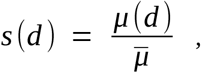

where 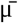 is the mean of μ(d) across all mapped DDIs. The selection score is the proportion of DDIs with s(d) < 1, hence those clustering more tightly than average. This selected ESM-2 with elementwise multiplication and a 128-dimensional latent space (Fig. S2).

For each DDI term with ≥30 mapped PPIs, we also scored local clustering purity in MAPPIE. For each member PPI we took its k=15 nearest neighbours and computed the fraction sharing the DDI term, averaged across members (sampling up to 500 members for each DDI). To compare common and rare domain pairs, we normalized by each terms background frequency to a fold-enrichment score by dividing the averaged purity by the background rate.

### UMAP projection

The selected 128-dimensional latent space was projected to 2D with UMAP (Euclidean metric, n_neighbors = 50, min_dist = 0.1, random_state = 42). The reducer was fit once unsupervised on the full reference set and reused unchanged for all figures and for placing query PPIs. All retrieval and enrichment use the 128-d latent space, and the UMAP is only used for visualisation.

### PPI projection

When both protein’s embeddings are already indexed in MAPPIE, we retrieve their embeddings directly; otherwise, we compute ESM-2 embeddings from user-supplied FASTA sequences or UniProt identifiers. We multiply the two protein vectors elementwise, standardize with the fitted scaler, and pass the result through the trained encoder to get the latent vector. We place this latent vector on the fixed UMAP using umap.transform. We further L2-normalize the latent vector for neighbor-search comparisons.

### Functional annotation

We annotated each PPI with the union of terms for both interacting proteins across nine sources retrieved from UniProt (38): GO biological process (BP), GO molecular function (MF), and GO cellular component (CC), KEGG and Reactome pathways, RHEA reactions, Pfam domains, InterPro families, and the potential domain–domain interactions prepared for the model selection. The term annotations cover a total of 17,723 GO terms (16, 17), 372 KEGG terms (18), 2,726 Reactome pathways (19), 6,933 PFAM (20) and 16,731 INTERPRO domains (21) and biochemical reactions from Rhea 4,754 (23) (Table S1).

Protein disorder fractions were taken from MobiDB’s consensus prediction-disorder-th50 (39).

### Neighbourhood enrichment

For a query PPI, we retrieve its *k* cosine nearest neighbours in the latent space using scikit-learn (40) *NearestNeighbors()* function and test each annotation term for over-representation against the full annotated PPI set as background using a hypergeometric test. Terms hit by at least one neighbour are tested, and *p*-values are corrected for multiple testing across this set with the Benjamini–Hochberg procedure (41), reporting significance for adjusted p < 0.05.

### Evaluation and Benchmarking

#### Functional coherence

To test whether neighbouring PPIs in latent space are grouped by shared function, we sampled 10,000 MAPPIE interactions and retrieved the *k* nearest latent neighbours at different (*k* = 10, 25, 50, 100, 200, 500) by cosine similarity for each PPI, excluding the query PPI itself. We then computed the enrichment of functional terms across all categories based on the union of neighbours, and reported the recall as the fraction of the query’s own terms recovered at adjusted p < 0.05. To test if the AE provides additional value, we repeated the identical procedure on the combined ESM-2 (13) embedding of each PPI, without further AE compression.

#### Complex coherence

Beyond single PPIs, we assessed whether MAPPIE also recovers functional terms annotated to protein complexes spanning a set of PPIs, retrieved from CORUM 5.3 (42). For each human protein complex, we checked all potential PPIs against the full HIPPIE (7) network and retained any protein complex with two or more PPIs annotated. This resulted in 302 human complexes with mappable subunits, curated GO terms and more than 2 subunits. For each of the complexes retained, we then computed the functional term enrichment on the union of the 100 nearest neighbours and tested the recall against the Functional Complex Group (FCG) annotation that CORUM (42) provides. We further report the resulting recall stratified by higher level functional root terms annotated in CORUM (42).

#### Comparison against homology and interactome baselines

Functional term annotation often relies on sequence homology and interactome association (9, 16, 25, 26). To test if MAPPIE’s performance depends on how well-characterized the functions of a query PPI already are, we stratified PPIs by their network degree. Specifically, we sampled up to 500 PPIs per degree bin (1–10, 10–25, 25–75, 75–100 and >100 partners) (Fig. S5). For each sampled PPI, we then took its first-degree neighbouring nodes from the HIPPIE network (7), defining *k* unique proteins. We then retrieved the same number of *k* proteins from the nearest neighbours in MAPPIE latent space, in the combined ESM-2 embedding (13), and by sequence identity, so that every method contributes an equally large set of unique proteins. For each of these sets we computed the enrichment of functional terms, reporting the recall as the fraction of the query PPI’s own terms recovered at adjusted p < 0.05. Sequence identity was computed with MMseqs2 easy-search default parameters (43), ranking candidates by the summed identity of the better-matching subunit pairing.

Across all four methods we excluded candidates sharing 30% or more identity with the query proteins (44). Without this, the sequence-identity baseline would retrieve near-duplicates of the query while the other methods could still draw on close homologs, biasing the comparison. For each set we computed functional-term enrichment and recorded recall as the fraction of the query PPIs own terms recovered at adjusted p < 0.05. To avoid circularity, GO annotations were restricted to experimentally supported evidence codes (16).

#### External network recovery

As a validation independent of HIPPIE (7), we checked how many of a query’s MAPPIE neighbours were also its neighbours in STRING v12 (6). For each MAPPIE PPI that is also present in STRING (159,152 pairs), we ranked all other PPIs by latent cosine similarity and skipped any that shared a protein with the query, since those would count as trivial recoveries. We then went down the ranked list and collected proteins until we had *k* unique ones (*k* = 10, 25, 50, 100, 200, 500).

We report the coverage as the fraction of this network among the *k* nearest MAPPIE proteins for different STRING confidence threshold (≥150, ≥400, ≥700, ≥900). As a baseline, we asked how much coverage we would expect by chance. For each protein in the STRING network (6), we used the hypergeometric distribution to compute how likely it is to appear in *k* randomly drawn PPIs, based on how many PPIs contain it, and averaged this across all network proteins. This accounts for hub proteins being more likely to come up by chance. Fold enrichment is the observed coverage divided by this expected coverage.

For each query, we walked its ranked neighbour list until k unique proteins were collected, counting a neighbour as a hit if either of its proteins was a STRING partner. Precision is the fraction of walked interactions that were hits, averaged across queries.

#### Dark protein interactome

We compute functional enrichment for PPIs that are currently poorly characterized, or so called dark. High-confidence interactions are taken from the BioPlex v3.0 293T network (5), using the published interaction-probability threshold of pInt ≥ 0.75, corresponding to 118,162 PPIs. Functional darkness is defined using the Unknome database, restricting to human proteins and classifying proteins with a knownness score ≤ 1.0 as dark (27). We intersect the BioPlex interaction network with these dark annotations and retain interactions in which both proteins are classified as dark. In total, the functional enrichment analysis is performed on 1,925 dark–dark PPIs spanning 1,236 proteins. We compute the functional enrichment of these interactions with MAPPIE using *k* = 100 neighbours and report adjusted p values < 0.05 as significant.

#### Web server

The MAPPIE web server is implemented in Dash (Python) (45) and hosted at https://cbdm-01.zdv.uni-mainz.de/∼mcihan/mappie/. Users can compute the functional enrichment for a candidate PPI or a set of PPIs, search the reference map by protein name or UniProt identifier, inspect individual interactions, and retrieve and visualise nearest neighbours in the latent space. Novel PPIs, defined from UniProt identifiers or user-supplied FASTA sequences, can be projected and analysed in the reference map.

## Acknowledgments

We thank Dr. Katja Luck for valuable input on the design of the evaluation framework. The authors gratefully acknowledge the computing time granted on the supercomputer MOGON 2/ MOGON KI at Johannes Gutenberg University Mainz (hpc.uni-mainz.de).

## Data availability

The code to train MAPPIE and to reproduce the evaluation benchmarks is deposited at https://github.com/mcihan0bioinf/mappie. The MAPPIE training datasets were derived from HIPPIE v2.3 (7), domain–domain interactions for model selection were obtained from 3did (22), and functional term annotations were retrieved from UniProt on 2026-01-24 (38). A publicly accessible web server is available at https://cbdm-01.zdv.uni-mainz.de/∼mcihan/mappie/.

## Fundings

The authors declare no specific funding for the conduct of this work. All research was carried out at Johannes Gutenberg University Mainz.

## Competing interest

The authors declare no conflict of interest with regard to this work.

## Notes

https://cbdm-01.zdv.uni-mainz.de/~mcihan/mappie/

### Competing Interest Statement

The authors have declared no competing interest.

