## Supplementary Information for "A map of human protein-protein interaction embeddings for functional discovery"

Figures

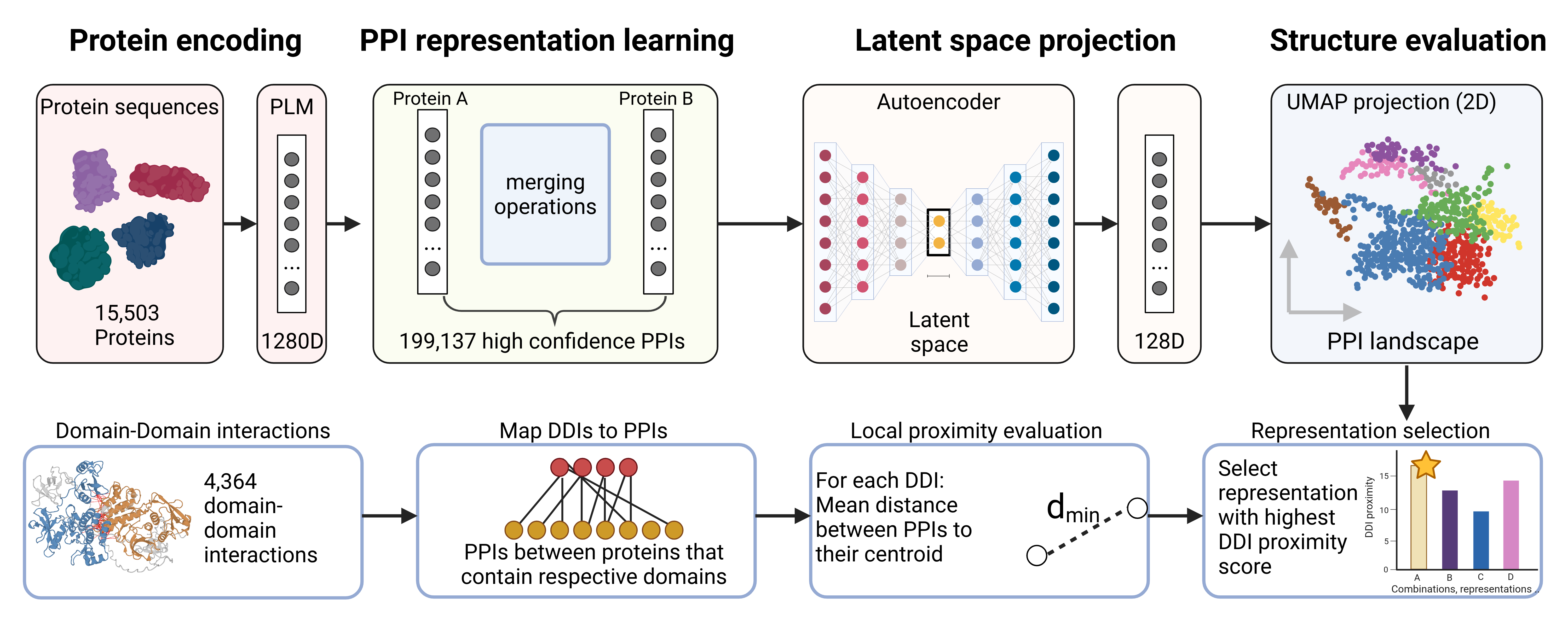

Fig. S1. MAPPIE architecture. Protein language model embeddings are combined per PPI, compressed by an autoencoder, and projected with UMAP into a two-dimensional PPI landscape, with domain–domain interactions used to select the best representation (Created with BioRender.com)

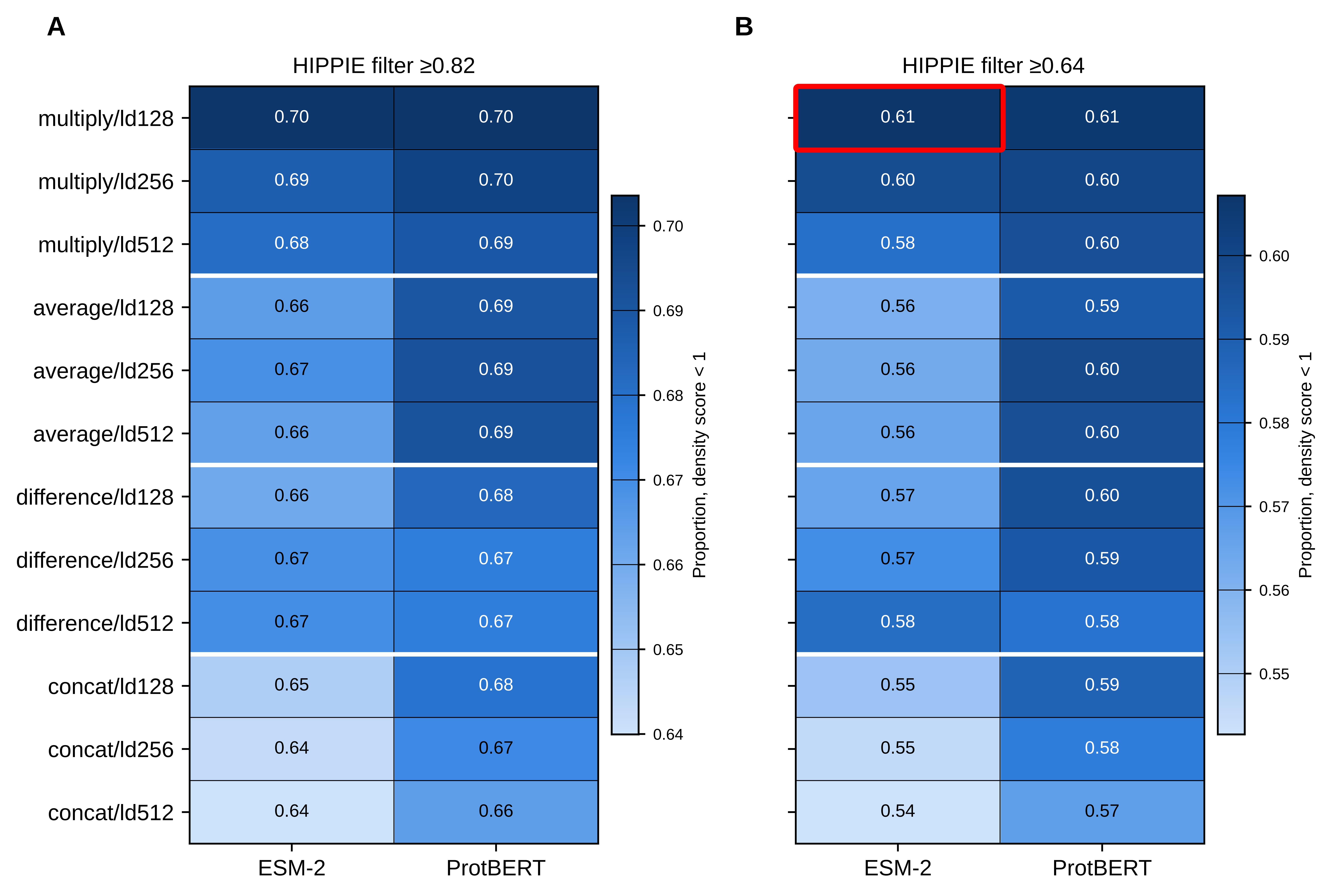

Fig. S2. Representation grid search for final model selection. For each combination of PLM embedding, merge operation and latent dimensionality, we show the proportion of DDIs whose mapped PPIs cluster more tightly than average. (A) High-confidence PPI subset. (B) Primary reference PPI set. The selected configuration (ESM-2, elementwise multiplication, 128 dimensions) is outlined in red.

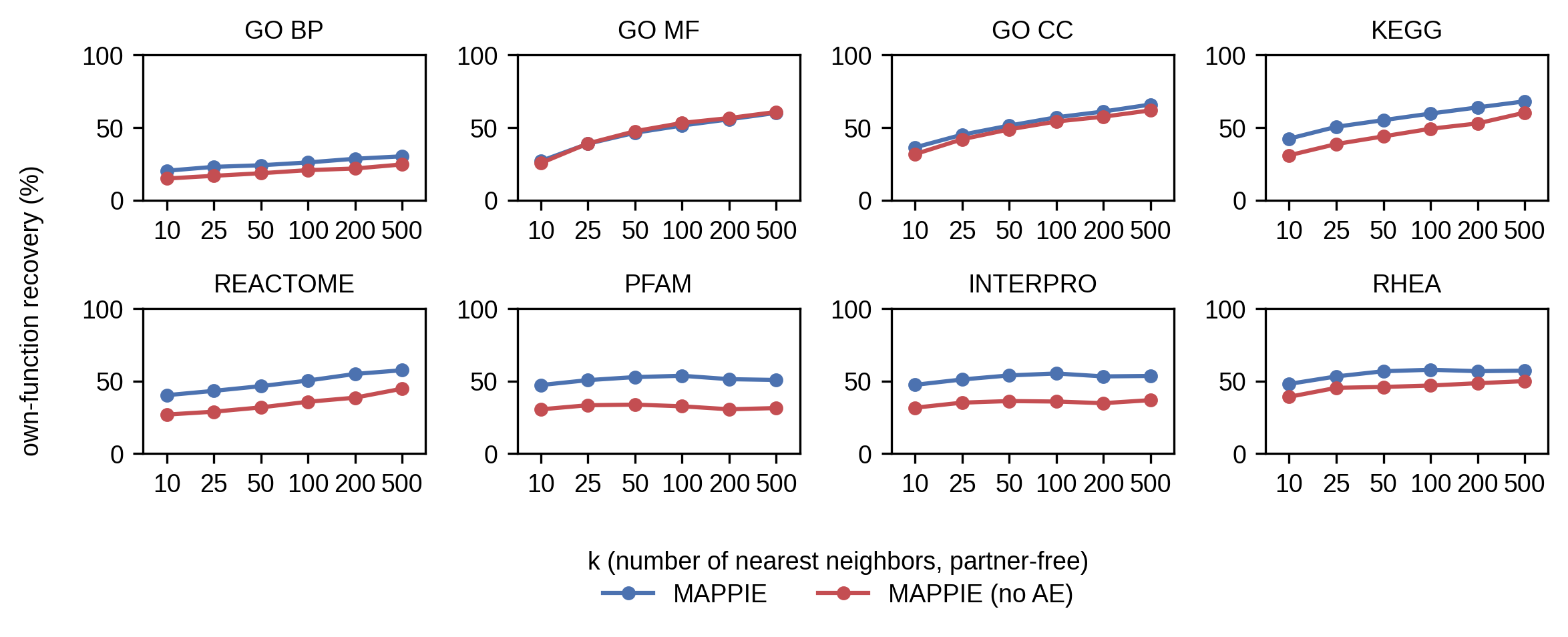

Fig. S3. Self recall excluding query proteins. Own-function recovery as a function of neighbourhood size for each annotation category, comparing the full MAPPIE model against a control using the interaction embedding without autoencoder compression. A total of 10,000 PPIs are sampled and for the enrichment analysis neighbouring PPIs that share one of the interacting query proteins are excluded.

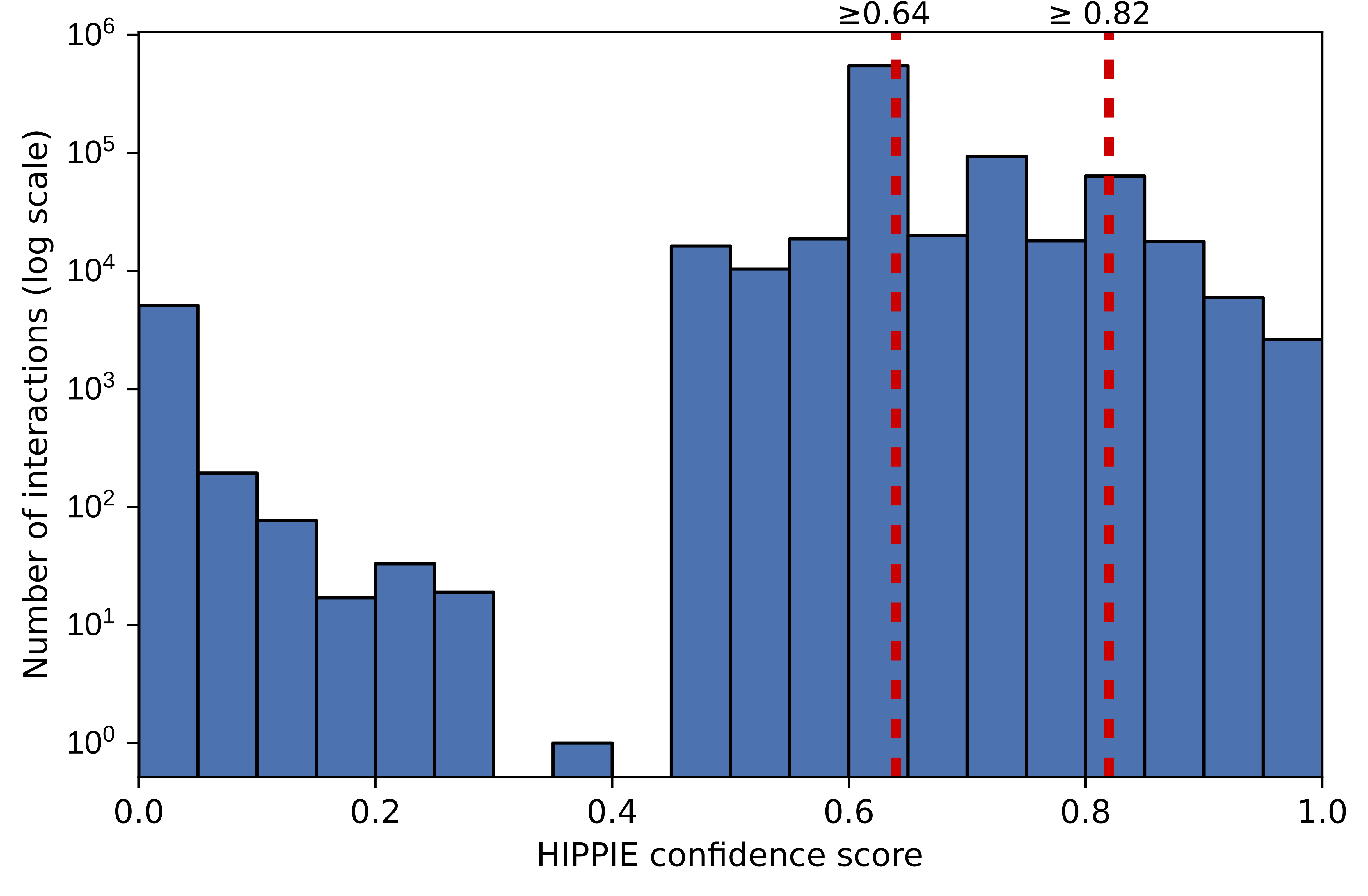

Fig. S4. Hippie confidence score distribution. Distribution of confidence scores across allinteractions in HIPPIE v2.3. Red dotted vertical lines mark the two thresholds used to define the MAPPIE training/reference sets: score ≥ 0.64 (199,137 PPIs, 15,503 proteins; 27% of all interactions) and the stricter high-confidence subset at score ≥ 0.82 (80,893 PPIs, top 10%).

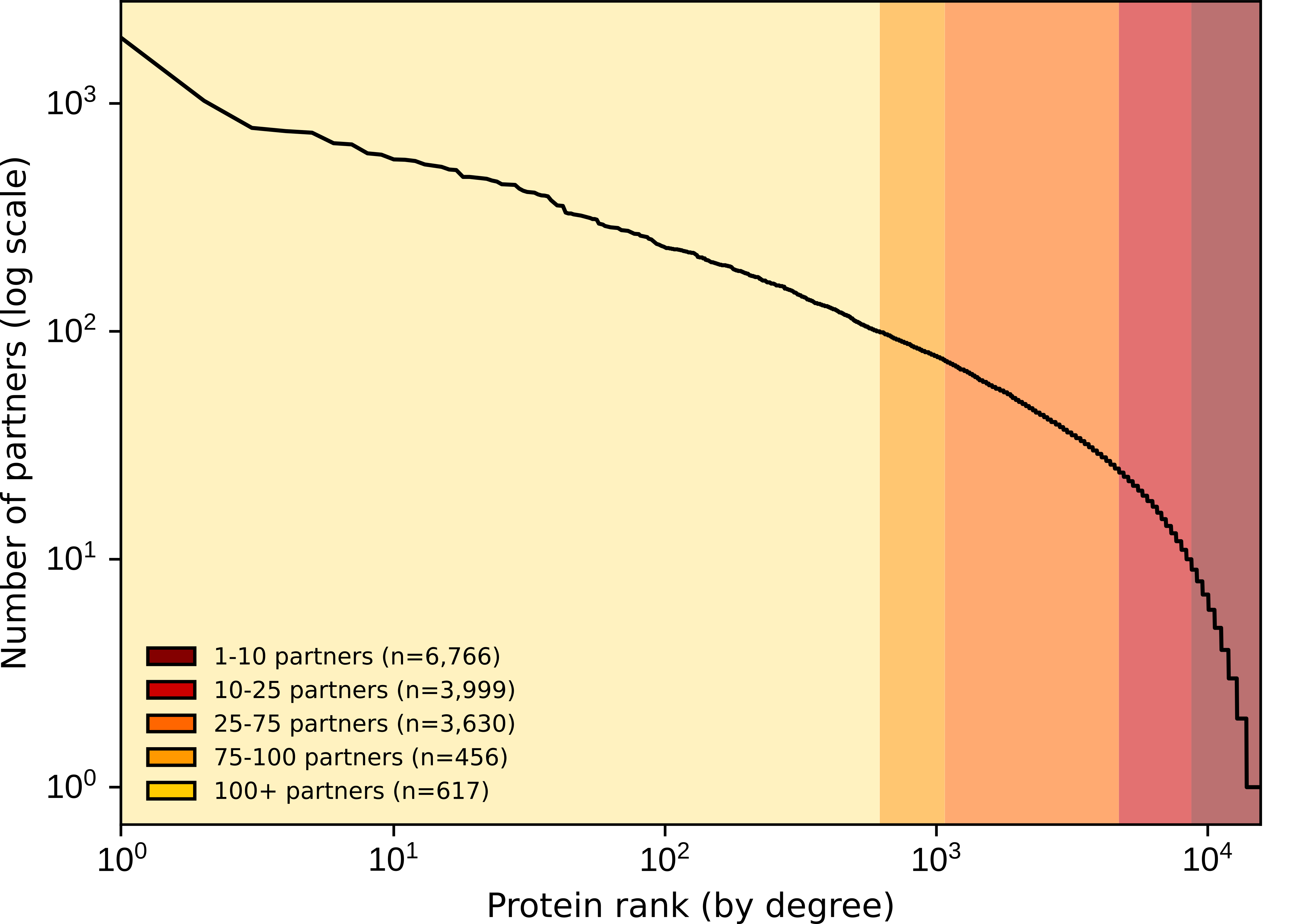

Fig. S5. Degree distribution of the reference interactome. Rank–frequency plot of the number of interaction partners per protein in the reference interactome (HIPPIE score ≥ 0.64).

Tables

Table S1. Overview of functional terms annotated in MAPPIE.

| Category | Terms | PPIs covered | Percentage PPIs | PPI-Terms |
| --- | --- | --- | --- | --- |
| GO BP | 11,575 | 197,820 | 99.30 | 4,944,383 |
| GO MF | 4,355 | 196,007 | 98.40 | 1,980,815 |
| GO CC | 1,793 | 198,536 | 99.70 | 2,593,401 |
| DDI | 6,374 | 27,355 | 13.70 | 48,081 |
| KEGG | 372 | 164,543 | 82.60 | 1,767,060 |
| REACTOME | 2,726 | 184,971 | 92.90 | 2,492,284 |
| PFAM | 6,933 | 198,190 | 99.50 | 766,064 |
| INTERPRO | 16,731 | 198,723 | 99.80 | 2,125,960 |
| RHEA | 4,754 | 76,322 | 38.30 | 202,917 |

Dataset S1 (separate file). Gene Ontology terms that are significantly enriched for the human dark protein interactome.
